# Automated cryo-lamella preparation for high-throughput *in-situ* structural biology

**DOI:** 10.1101/797506

**Authors:** Genevieve Buckley, Gediminas Gervinskas, Cyntia Taveneau, Hari Venugopal, James C. Whisstock, Alex de Marco

## Abstract

Cryo-transmission electron tomography (cryo-ET) in association with cryo-focused ion beam (cryo-FIB) milling enables structural biology studies to be performed directly within the cellular environment. Cryo-preserved cells are milled and a lamella with a thickness of 200-300 nm provides an electron transparent window suitable for cryo-ET imaging. Cryo-FIB milling is an effective method, but it is a tedious and time-consuming process, which typically results in ~10 lamellae per day. Here, we introduce an automated method to reproducibly prepare cryo-lamellae on a grid and reduce the amount of human supervision. We tested the routine on cryo-preserved *Saccharomyces cerevisiae* and demonstrate that this method allows an increased throughput, achieving a rate of 5 lamellae/hour without the need to supervise the FIB milling. We demonstrate that the quality of the lamellae is consistent throughout the preparation and their compatibility with cryo-ET analyses.

## Introduction

Today, the best method to image the cellular environment in its native state and obtain structural information is Cryo-Electron Tomography (cryo-ET) (Beck & Baumeister, 2016). Biological samples are vitrified by fast freezing at cryogenic temperature (80 – 120 Kelvin) and then imaged using a cryo-Transmission Electron Microscope (TEM) following a tomographic acquisition scheme. Cryogenic fixation is instantaneous and it helps to reduce the effects of radiation damage during data acquisition (Dubochet et al., 1988). This process ensures that all the water contained in the sample is vitreous, therefore solid and capable of withstanding the high-vacuum environment of TEMs while not diffracting the electron beam. As a general guideline, the maximum allowed sample thickness is ~300nm, since most cryo-TEM operate at 300 keV where an electron statistically undergoes a single elastic scattering event in ~250 nm of water (elastic mean free path) (Holtz, Yu, Gao, Abruña, & Muller, 2013). Thicker samples provide higher chances of inelastic scattering, which in non-crystalline samples cannot be interpreted as useable information. Since cells are typically multiple microns thick, the inspection is limited to the thinnest regions such as filopodia (Carlson et al., 2010; Stauffer et al., 2014; Weber, Wojtynek, & Medalia, 2019). The first approach that made it possible to inspect the inside of a cryo-preserved cell through cryo-ET was cryo-sectioning of vitreous sections (CEMOVIS) (Al-Amoudi et al., 2004). CEMOVIS has the advantage of potentially allowing serial-section tomography with minimal losses throughout the cell, but it is extremely difficult to perform and suffers from mechanical deformations and is prone to surface contamination (Al-Amoudi, Studer, & Dubochet, 2005). Alternatively, using an extremely common approach in semi-conductor sciences, it is possible to isolate a thin lamella of material using a dual-beam microscope which utilizes a Scanning Electron Microscope (SEM) for imaging and a Focussed Ion Beam (FIB) for machining any sample with nanometre resolution (Marko, Hsieh, Schalek, Frank, & Mannella, 2007; Rigort et al., 2012; Schaffer et al., 2015; Schaffer et al., 2017). Cryo-FIB milling has been shown to be a highly effective method for preparing cellular sections for cryo-electron tomography (Rigort et al., 2012; Schaffer et al., 2015; Schaffer et al., 2017; Villa, Schaffer, Plitzko, & Baumeister, 2013), but it is time-consuming since the resolution and the milling speed are inversely proportional.

In contrast to industrial FIB applications (e.g. semi-conductors), all existing procedures for cryo-lamellae preparation are performed manually due to the heterogeneity and dose-sensitivity of biological specimens. The immediate effect is that highly trained users are performing an extremely repetitive task. This not only results in an elevated cost, but it also restricts production to a throughput of 5-10 lamellae per day (Danev, Yanagisawa, & Kikkawa, 2019; Koning, Koster, & Sharp, 2018; Wolff et al., 2019). Further, in most cases, cryo-FIB microscopes are only used for one session/day while users are onsite, therefore to boost the productivity night shifts are the only way forward. Cryo-ET acquisition can already run unsupervised and regardless of the acquisition scheme used it is possible to obtain at least 20 tomograms over 24 hours. Data collection schemes optimized for high resolution will require ~60 min of acquisition time per tomogram (Hagen, Wan, & Briggs, 2017), while fast tomography workflows will allow for acquisition times down to 5 minutes (Chreifi, Chen, Metskas, Kaplan, & Jensen, 2019; Eisenstein, Danev, & Pilhofer, 2019). Moving beyond the bare throughput and instrument usage, since FIB milling is a manual and iterative process the reproducibility scales with the user experience. Recognising the need for reproducible sample preparation, several workflows and best practice procedures have been developed (Hsieh, Schmelzer, Kishchenko, Wagenknecht, & Marko, 2014; Medeiros et al., 2018; Schaffer et al., 2015), but the initial steep learning curve still represent a limitation especially for laboratories where in house expertise has not yet been established.

One way to counteract the low throughput of cryo-lamella preparation consists of targeting a region based on the known presence of an interesting event or structure. In order to achieve this level of efficiency, the typical approach uses correlative light and electron microscopy techniques which, when properly performed, prevents from imaging areas that do not contain useful information for a specific study (Arnold et al., 2016; Gorelick et al., 2019). Regardless of the correlative microscopy approach in use, the mismatch between the throughput of cryo-ET and the preparation of lamellae is evident. Here we introduce an automated and reproducible method for on-grid cryo-lamella preparation for vitrified cell. We demonstrate an increased throughput capable of reproducibly producing ~5 lamellae/hour. Since the presented protocol can run unsupervised, the learning phase for new users will be shorter than the current situation. Further, with the appropriate cryo-stage, this protocol opens the possibility for a 24/7 cryo-lamellae preparation.

## Materials and methods

### Cell culture and sample cryo-fixation

*Saccharomyces cerevisiae* was cultured in YPD at 30 degrees, cells were harvested when an optical density of 0.6 was reached. Cells were washed twice in PBS and re-suspended in half of the initial volume. 5 μL of cell-containing solution was deposited on a Cu-300 R2/2 grid (Quantifoil™) which had been previously glow discharged for 30 seconds using a Pelco EasyGlow™. Excess solution was removed through manual blotting and cells were vitrified by plunge freezing in a liquid 60/40 Ethane/propane mix and stored in liquid nitrogen.

### Cryo-FIB milling

All tests presented here were conducted on a ThermoFisher Helios Ux G4 DualBeam™ equipped with a Leica cryo-stage and a VCT500 cryo-transfer system. Grids were clipped in autogrid cartridges prior to loading into the cryo-FIB.

### Cryo-FIB control and scripting

All manual operations were conducted through the ThermoFisher Xt UI™, while batch cryo-FIB operations were performed using Python 3.6 through the communication API ThermoFisher Autoscript 4.1™ (which is required to control the microscope). Scientific python packages used include numpy, scipy, matplotlib, scikit-image, click, pyyaml and pytest (Hunter, 2007; Oliphant, 2007; S. van der Walt, Colbert, & Varoquaux, 2011; Stéfan van der Walt et al., 2014).All original scripts used in this procedure are installable via the pip package installer for Python. All the source code and the instructions can be downloaded from the laboratory webpage (www.github.com/DeMarcoLab/autolamella).

### Cryo-TEM

Cryo-electron tomograms and micrographs were collected on a ThermoFisher Titan Krios G3 equipped with a post-GIF Gatan K2™ DDE camera. Images were collected in EFTEM mode following a low-dose acquisition scheme. The microscope setup included a 50 μm condenser aperture and a 70 μm objective aperture. All tomograms were imaged from −60 to +60 deg with 3 deg increment. The total dose on the tilt-series was 98.4 e^−^/Å^2^, with a dose-rate of 4e^−^/pixel/sec and tilt micrographs were collected at a nominal magnification of 42000x resulting in a final pixel-size of 3.61 Å. Low magnification overviews were collected at a nominal magnification of 2500x in LM mode, resulting in a final pixel-size of 20 nm.

### Tomography reconstruction

Tilt-series were aligned using patch tracking and back-projected using Etomo from the IMOD suite (Mastronarde, 1997). Tomogram visualization and filtering occurred through IMOD or UCSF Chimera (Pettersen et al., 2004).

### Image display and figure preparation

All figures have been prepared using Adobe Illustrator. Cropping, filtering and contrast adjustments have been performed using IMOD or Fiji. Videos have been made using Fiji (Schindelin et al., 2012).

## Results and discussion

Any TEM sample preparation procedure based on FIB milling requires the following operations: changing the milling current and/or energy, changing the imaging parameters, stage movements for sample realignment, beam-shift for drift compensation and setup/positioning of the milling patterns. On average, every milling step can require time spanning from a few seconds to multiple minutes and the sequence is extremely repetitive and predictable. In this situation, even the most diligent operator will incur a significant time-waste as a consequence of short delays at the start of every step and due to the time required to perform manual realignment.

Here, we analysed the preparation workflow and reduced its complexity to the 4 steps: (i) identification of the regions of interest; (ii) opening the lamellae (trenching); (iii) thinning and (iv) polishing the lamellae. Cryo-FIB milling occurs in a high-vacuum environment where the pressure rages from 2e-6 to 5e-7 mBar. Under these conditions the atmosphere around the sample is not completely depleted from water, therefore, the slow growth of a crystalline ice layer on the sample surface is expected over time. Typical contaminations rates measured as crystalline water deposition are specified by the instrument manufacturers to be in the order of 5 – 20 nm/hour. In addition, the material that is sputtered while milling can redeposit on neighbouring areas, therefore contaminating finished lamellae. To limit contamination, we concluded that the best course of action is to run the lamella preparation protocol in 3 or more stages. Accordingly, we defined a default protocol that includes 3 stages, first the routine loops through all the regions of interest and open the trenches, then lamellae are thinned to ~500 nm and last they are polished to the desired thickness (100-300 nm). Following this protocol, the bulk material removal, which also represents the most time-consuming part is carried out over the first two steps, while the polishing step is performed over the last few minutes prior to unloading the sample. The default number of stages for the preparation of the lamellae is three, but users can add or remove stages if the number of conditions required to achieve good lamellae is different on other samples. The presented routine includes stage movement, beam current changes and, most importantly, it requires time. Accordingly, there will be drift and relocation errors to be accounted for. Our proposed protocol follows a low-dose approach similar to the one performed in cryo-ET, where a sacrificial area containing a fiducial marker is used for tracking and focussing, whereas the sample area is imaged only once at low-dose and magnification to define the location of the region of interest.

**Figure 1:**
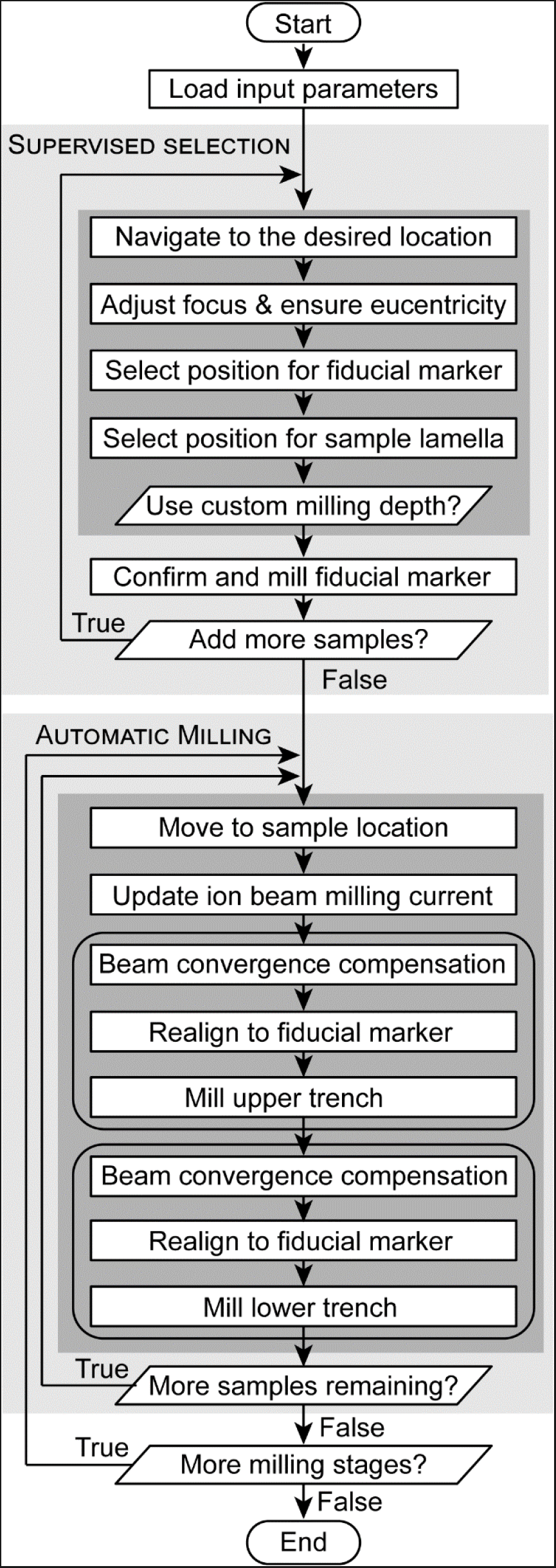
Flowchart of the automated cryo-lamella process. First, a configuration file containing all the required parameters is loaded (e.g. lamella size, currents, etc.). The process can be divided into two sections, a human supervised section followed by the automated lamella preparation across all selected locations. The supervised section requires users to interactively select the locations. The general navigation can be performed through the ThermoFisher Xt UI™ or ThermoFisher MAPS™ (where available). In order to achieve an optimal performance, the user must adjust the focus and ensure the sample stage is at the coincidence point between the FIB and SEM beams. The position of the fiducial marker (used for image realignment) and the lamella can be defined anywhere within the field of view. This section is performed as many times as the number of lamellae required. Once all sites of interest have been selected the automated section starts. Here, automated ion beam milling is performed for all selected samples. First, the initial trenches are opened for every sample (typically using a higher milling current). Then the FIB milling current is reduced and every sample is further thinned until the final polishing step. Every time the sample stage is moved or the FIB milling current is changed, the sample position is checked using the fiducial marker and correction is done through beam-shift. The compensation for the beam convergence can be specified in the input configuration file and can be changed for each milling stage.

Figure 1 shows a flowchart diagram of the program logic. We minimized the required user interaction: the user provides a configuration file when launching the program (an example file is provided with the package) which contains all the conditions required for the lamella preparation. The parameters defined in the configuration file are fixed for each batch job and include the milling and imaging currents, the depths, the field of view size, as well as the fiducial and the lamella sizes. The milling depth for lamellae and fiducials are set in the configuration file, but, where required, they can be set independently for each location. This decision was made to provide a good compromise between the flexibility and setup time. Lamellae locations are interactively selected together with the location of the tracking area. In order to improve reproducibility, a small cross-shaped fiducial marker is FIB milled in the centre of the tracking area. The fiducial marker assists image re-alignment after stage movements and shifts due to changes in the ion beam current. The image containing the fiducial marker is updated after every re-alignment to take care of eventual degradations that might occur in response to imaging. In principle, this area can be used to correct against drift during the milling procedure, but we found that on two systems with different cryo-stages (both ThermoFisher Helios G4) this option is not needed given the short time that each milling stage requires. Since the sample thickness can vary substantially across locations, we kept the possibility to adjust the desired FIB milling depth individually for each lamella and fiducial. This setup procedure is repeated for as many locations as the user wishes to mill and the only constraint here is the time required for milling. The user is guided step-by-step throughout the setup phase (for examples through the workflow see supplementary figures 3–6). The tracking area is also used to re-align the lamella position when compensating for the beam convergence (Schaffer et al., 2017). For this option, users are able to define the amount of over/under-tilt to apply in relation to the current and instrument in use.

Since the tracking procedure follows a low-dose acquisition scheme, the site where the lamella will be prepared is going to be imaged only once using the ion beam and twice using the SEM (optional). In order to help the users during the protocol optimization for each sample and for documentation purposes, we added the option to acquire full-field ion beam images at each stage of the process, but in general, this option would not be used so the sample conditions can remain pristine. The ion beam image that is acquired at the beginning of the process to define the lamella position and the tracking area generally will have a low resolution and fast dwell time to minimize the dose applied to the sample. In our experiments, we have found that a field of view of 50 μm acquired with a pixel size of 32nm provides a good image quality when using 10-30 pA of current with dwell times comprised between 50 and 300 nsec. Under these conditions, the dose is extremely low but the alignment precision is limited by the pixel size. Accordingly, we have set the possibility of acquiring the tracking images (restricted to the sacrificial area) with a smaller pixel size (e.g. 8 or 16 nm) and dose, therefore making the alignment process significantly more precise. Further, in order to increase the flexibility in case a user needs to image samples that do not align efficiently with the provided algorithm, we set up the code structure to allow custom frequency filtering and real space masks to be applied, or even the replacement of the alignment function with another algorithm.

In figure 2 we show a gallery through the lamella preparation procedure. The cross mark visible in the image represents the centre of the tracking area and has a size of 6×6 μm. In order to respond to the variability of the sample topology, we made sure that the position of the fiducial and the lamella can be selected and do not have a strict relative position. The only limits set are to prevent the milling areas to overlap between fiducial and lamella as well as to prevent any of the two areas to be outside the field of view.

**Figure 2:**
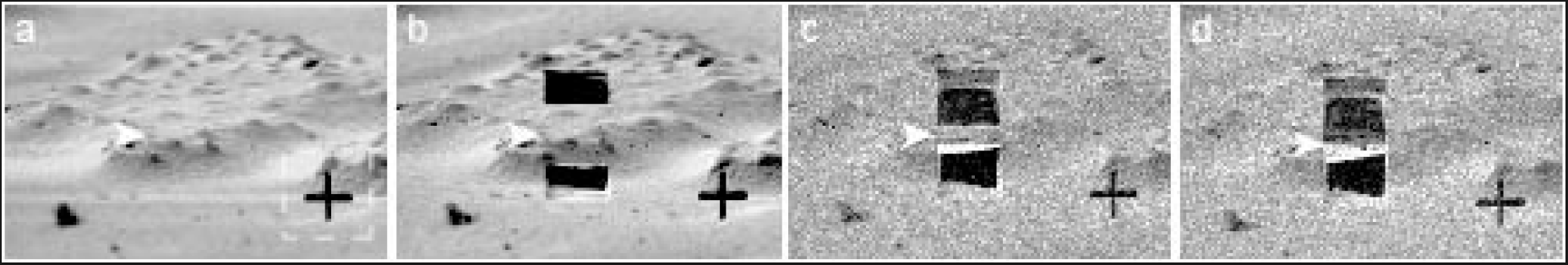
Ion beam images of a single lamella during the automated FIB milling. (a) A cross-shaped fiducial marker is created for later image realignment, reduced area field of view used for realignment shown by dashed white box; (b) trenches are milled above and below the lamella using a current of 2.4 nA; (c) the lamella is thinned further using an intermediate ion beam current, 75 pA; (d) the completed lamella after final ion beam milling at low current, 26 pA. White arrowheads indicate the position of the finished lamella; the scale is indicated by 6×6 μm cross-shaped fiducial marker.

We tested the robustness of the procedure by setting up batch preparation jobs with multiple (3-5) lamellae. Our results indicate high reproducibility throughout each job with lamellae thicknesses always comprised between 210 nm and 250 nm. The throughput was measured to be 4 to 5 lamellae/hour depending on the sample thickness. In supplementary figure 1, we show a gallery comprising 5 lamellae which have been prepared in 47 minutes from the beginning of the batch procedure. The setup phase (selecting sites and defining tracking markers) for that job required approximately 12 minutes. In order to ensure that the quality of the samples is not compromised, we inspected all lamellae through cryo-TEM (see figure 3 and supplementary figure 2), and imaged a subset via cryo-ET (figure 4 and supplementary video), therefore ensuring the suitability for downstream analyses. The level of curtaining is comparable to what has previously been reported from manual preparation. Although in this case, we have noticed that the length of the lamella directly correlates with the level of curtaining. This observation correlates with the fact that we applied the same ion dose when preparing all lamellae (supplementary figure 2).

**Figure 3:**
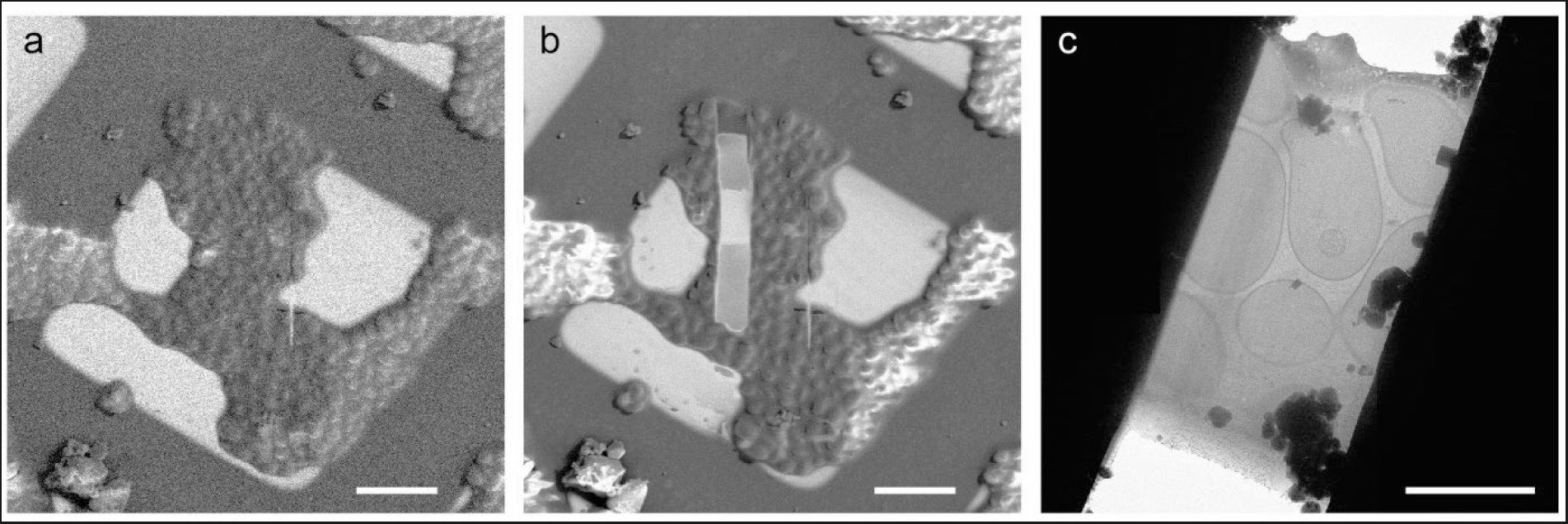
View of an exemplary lamella from cryo-SEM and low magnification cryo-TEM. (a) shows a cryo-SEM view of a region before the lamella preparation; (b) shows the same region after sample preparation. Both SEM images have been acquired as part of the automated procedure. (c) shows a low magnification cryo-TEM overview of the same lamella displayed in a and b. The surface quality is comparable to what has previously been reported from manual preparation. Scale-bars in (a) and (b) are 20 μm, while in (c) the scale bar 5 μm.

**Figure 4:**
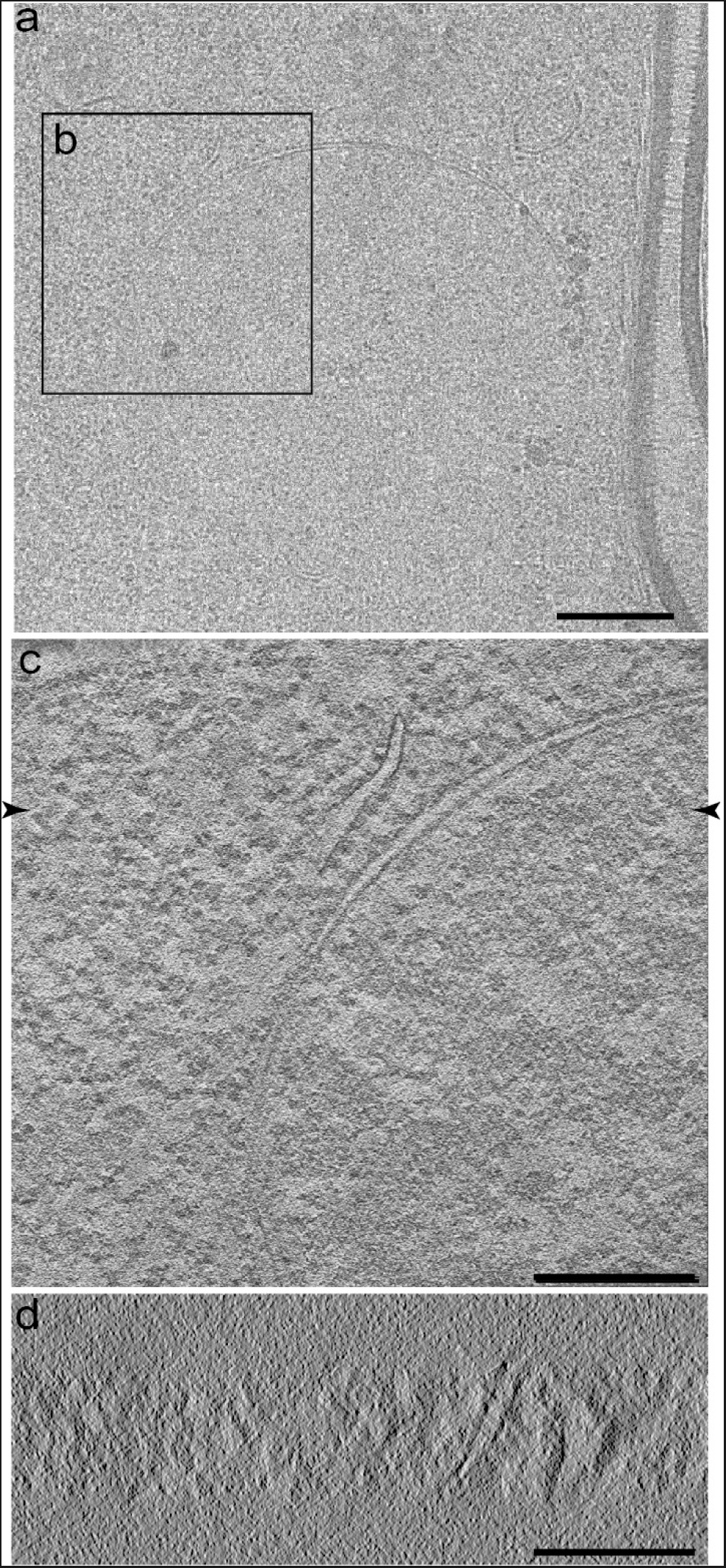
Cryo-TEM and cryo-ET performed on the same lamella shown in figure 2. Panel (a) shows a mid-magnification (6500x) micrograph of the lamella. Inset (b) in panel (a) corresponds to the XY slice from the tomogram collected and displayed in (c). Panel (d) shows a XZ slice through the tomogram. The arrowheads in (c) show the position of the slice in displayed in (d). Scale-bars are 500 nm in (a) and 200 nm in (c) and (d).

Our results show that by automating the lamella preparation it is possible to prepare uniform lamellae with the desired thickness at a rate which is far greater than what could be reproducibly achieved with manual operation. The average preparation time for 1 lamella is ~9 minutes, excluding sample loading, initial alignment and selection of the regions of interest. From experience, users will require about 1 hour for stage cooling and sample loading, then Pt deposition and general imaging of the samples for navigation will require ~10-15 min. Then, when the procedure starts, the basic navigation is conducted through the microscope UI and the user must ensure that when a region of interest is selected the sample is at the coincidence height. This is important because the overview images acquired with the SEM before and at the end of the milling procedure can provide immediate feedback on the quality of each lamella before going to the cryo-TEM. If the stage is not at the correct height the SEM images will not include the lamella site. Further, if the user chooses to compensate for the beam convergence (optional), the stage tilt will change by 1-4 deg. If the stage is not at the correct height the region of interest will move outside the field of view. Our experience shows that the user supervised part of the procedure which includes the selection of the regions of interest and the preparation of the fiducial requires ~2-3 minutes/site. Accordingly, if a user loads 2 grids and selects 20 sites/grid the preparation time from stage cooling to the completion of 40 lamellae would be between 10 and 11 h, with the polishing step conducted through the last 1.5 h.

The typical duration of a cryo-FIB session is limited by the liquid nitrogen supply (ThermoFisher and Quorum cryo-stages can be kept cold for ~11 h, while Leica cryo-stages have the possibility to refill the liquid nitrogen dewar and provide sessions that are longer than 24 h). The automation and the fact that the procedure runs mostly unsupervised ensures that for ~8 h users do not need to be on-site. Accordingly, this opens up the possibility to prepare cryo-lamellae with a 24 h continuous operation of the microscope and therefore lead to ~80 or more lamellae/day. Considering the latest developments in regards to fast cryo-ET acquisition it will be possible to match this throughput on the TEM (Chreifi et al., 2019). Even in the situation where the above-described throughput is not required, a protocol providing up to 40 lamellae during a daytime session will shorten the user queue around cryo-FIBs, therefore, allowing more projects to be conducted in facilities where the availability of instrumentation is limited in relation to the number of users.

## Acknowledgements

We acknowledge the support and expertise of the Clive and Vera Ramaciotti Platform for Structural Cryo-Electron Microscopy (Monash University, Clayton). GB, CT, JCW and AdM acknowledge the support of an Australian Research Council Laureate Fellowship and the ARC Centre of Excellence in Advanced Molecular Imaging. JCW further acknowledges the support of the National Health and Medical Research Council of Australia Senior Principal Research Fellowship and Program grant. We also thank Sergey Gorelick for his contribution to the graphical abstract.

---

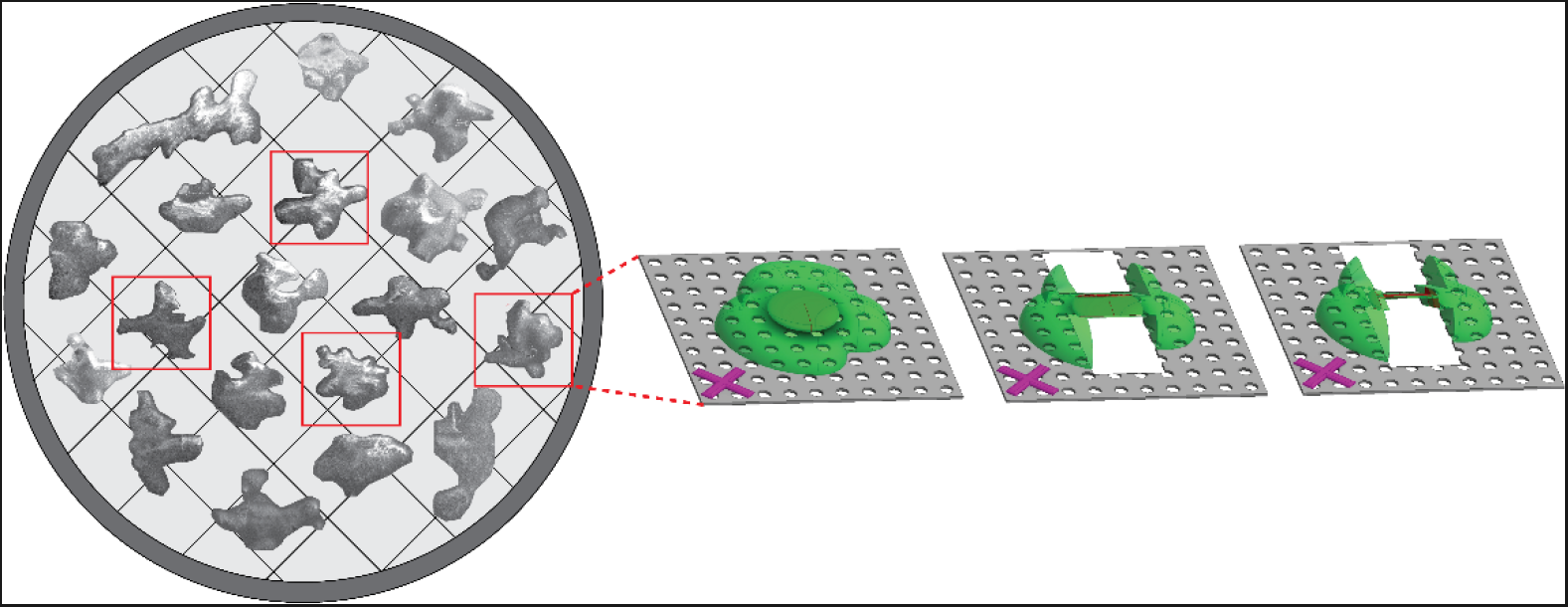
Graphical abstract.

## Supplementary Information

**Supplementary figure 1:**
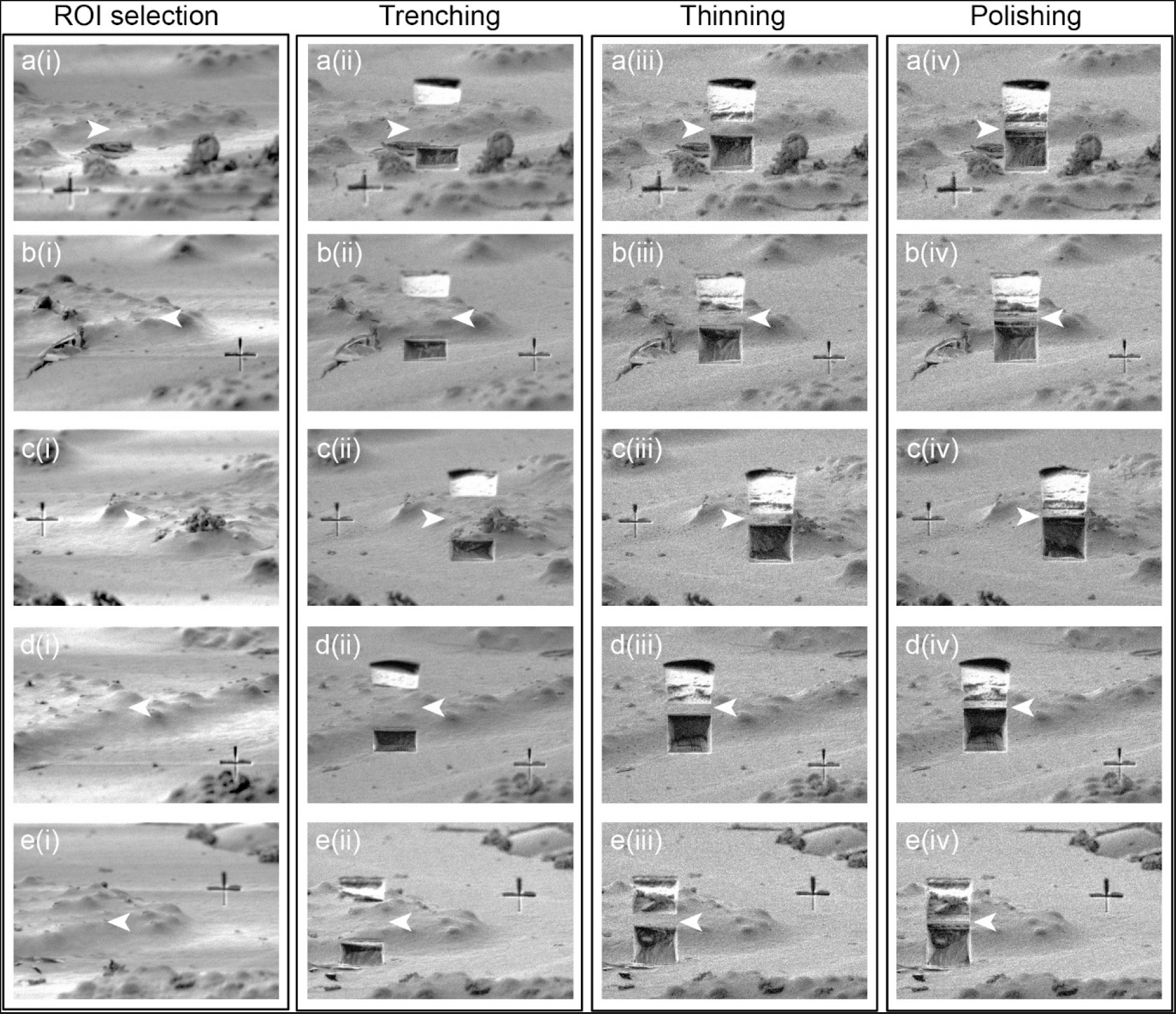
Ion beam images acquired throughout a single batch job consisting of 5 lamellae. The total duration of the batch job was 47 minutes, plus 12 minutes required for the initial (manual) selection of sample locations by the user. Here, each lamella location is displayed in a single row (marked a, b, c, d, e) and white arrowheads indicate the position of the finished lamella. The cross-shaped fiducial marker acts as scale-bar, with dimensions of 6×6 μm. Columns from left to right show: (i) the fiducial marker is created; (ii) trenches opened with high ion beam current; (iii) further thinning with an intermediate ion beam current and (iv) the completed lamella after final polishing with low ion beam current. Ion beam milling currents for these stages were 2.4 nA, 75 pA, and 26 pA, respectively. The milling procedure runs per column, therefore, each FIB milling stage is completed for every lamella location before proceeding to the next milling stage.

**Supplementary figure 2:**
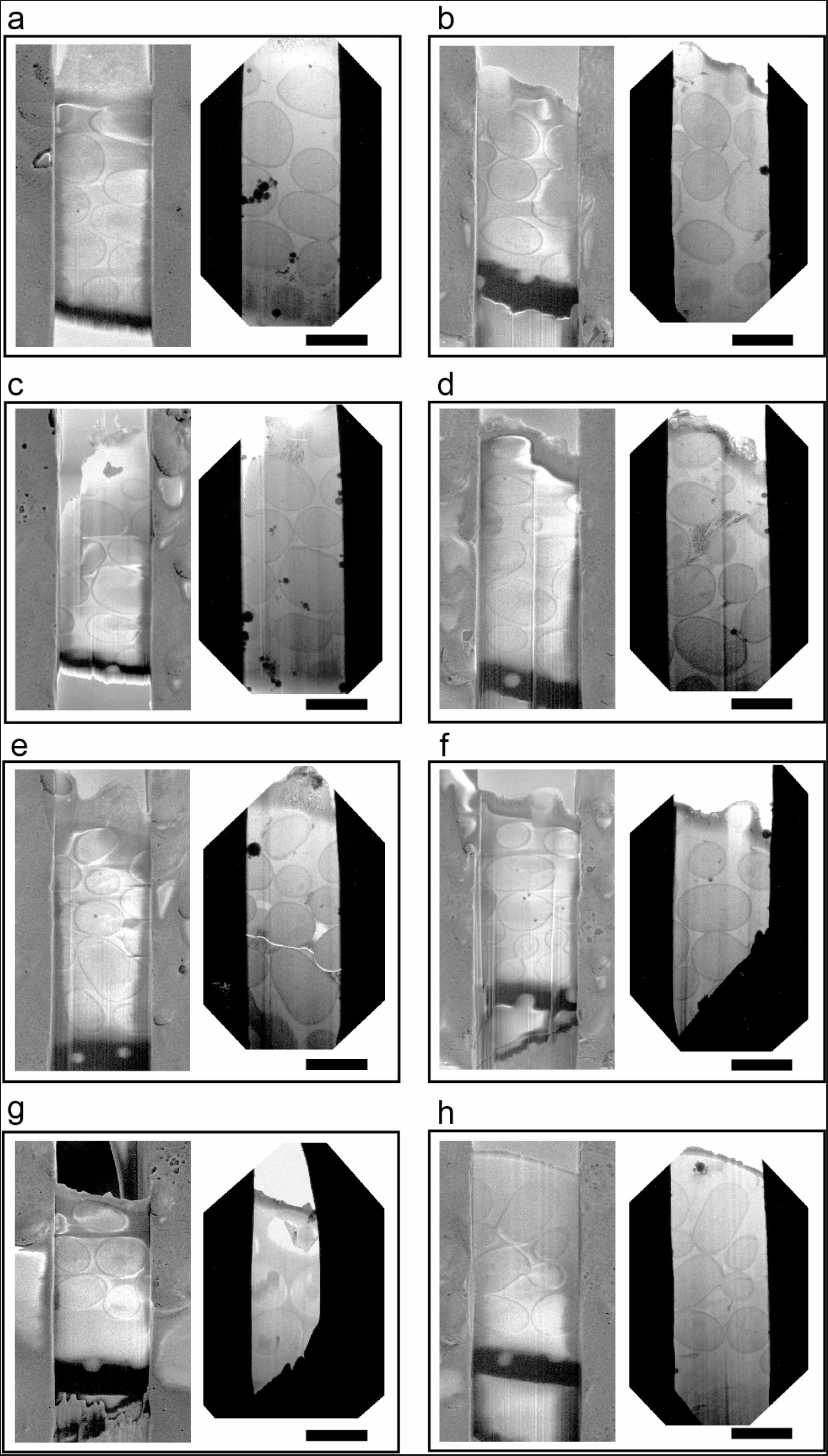
Gallery of 8 lamellae prepared using the procedure described in this article, illustrating quality and pitfalls. Each panel shows on the left a low voltage cryo-SEM image of the lamella surface at the end of the preparation procedure. On the right within each panel there is a low mag cryo-TEM micrograph for each lamella. The images show that the quality is comparable to current preparation approaches. In this gallery, we show potential pitfalls of batch preparation which can be easily resolved if the user is aware and careful. The level of curtaining changes in response to lamella length and surface contamination, if the sample thickness is variable the user should select more generous depth for the polishing step to ensure the surface finish is optimal. (a, b) show proper depth selection, while (c–h) show different behaviours when under-polishing). Panels (f) and (g) show that the size of the trenches was not appropriate for the sample thickness, as visible from the cryo-TEM images part of those lamellae is not electron transparent, indicating that there is bulk material underneath the lamella (also visible from the cryo-SEM). Scale-bars are 5 μm.

**Supplementary figure 3:**
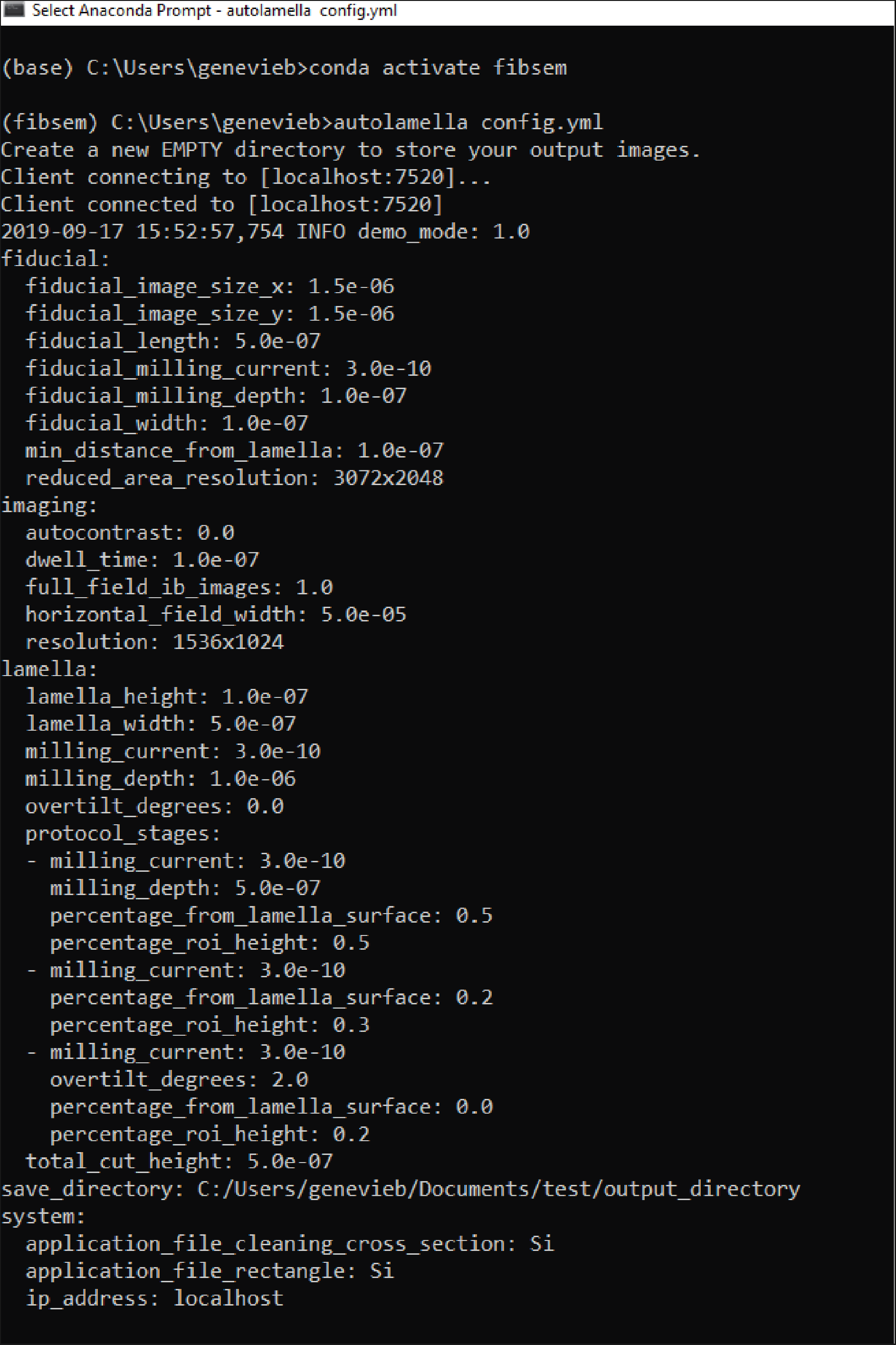
Screenshot of the terminal as the user launches the ‘autolamella’ program. The first section of the output consists of a summary of the parameters provided in the configuration file, here named config.yml. The idea is to provide immediate feedback that the file and the parameters selected are the correct ones. This screenshot demonstration has been produced using the Autoscript simulation mode.

**Supplementary figure 4:**
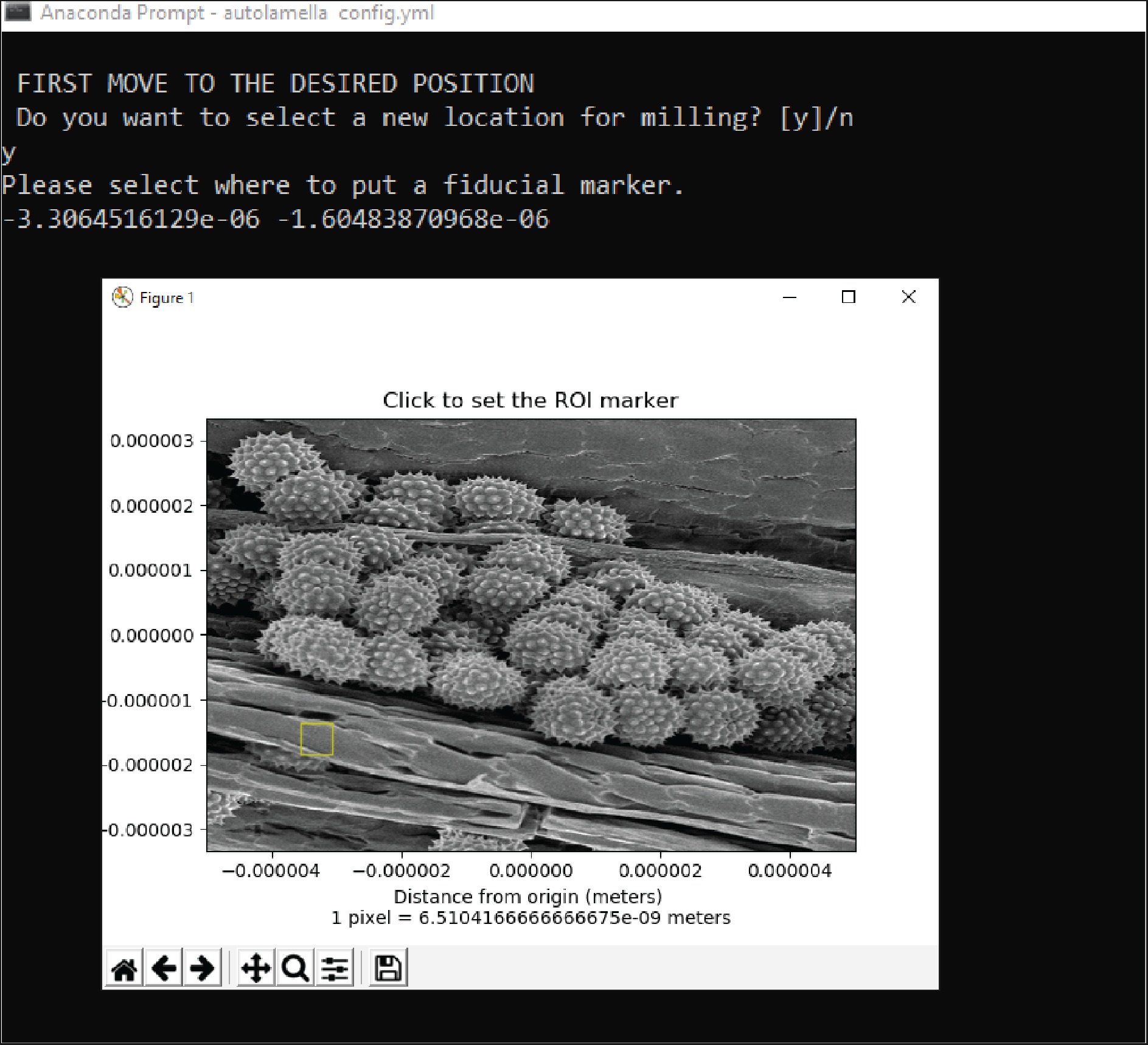
Screenshot of the interactive selection of the tracking area position. The user finds a suitable location for a lamella interactively using the ThermoFisher Xt UI™. Once the new location has been identified, the user replies yes to the question in the command prompt, “Do you want to select a new location for milling?”. The current ion beam image is then displayed in a pop-up window, and the user clicks to select the position for the fiducial marker. The fiducial marker position is indicated with a yellow box, and this position can be adjusted before closing the pop-up window. Closing the window leads to continuing the process. Screenshots have been produced using Autoscript simulation mode.

**Supplementary figure 5:**
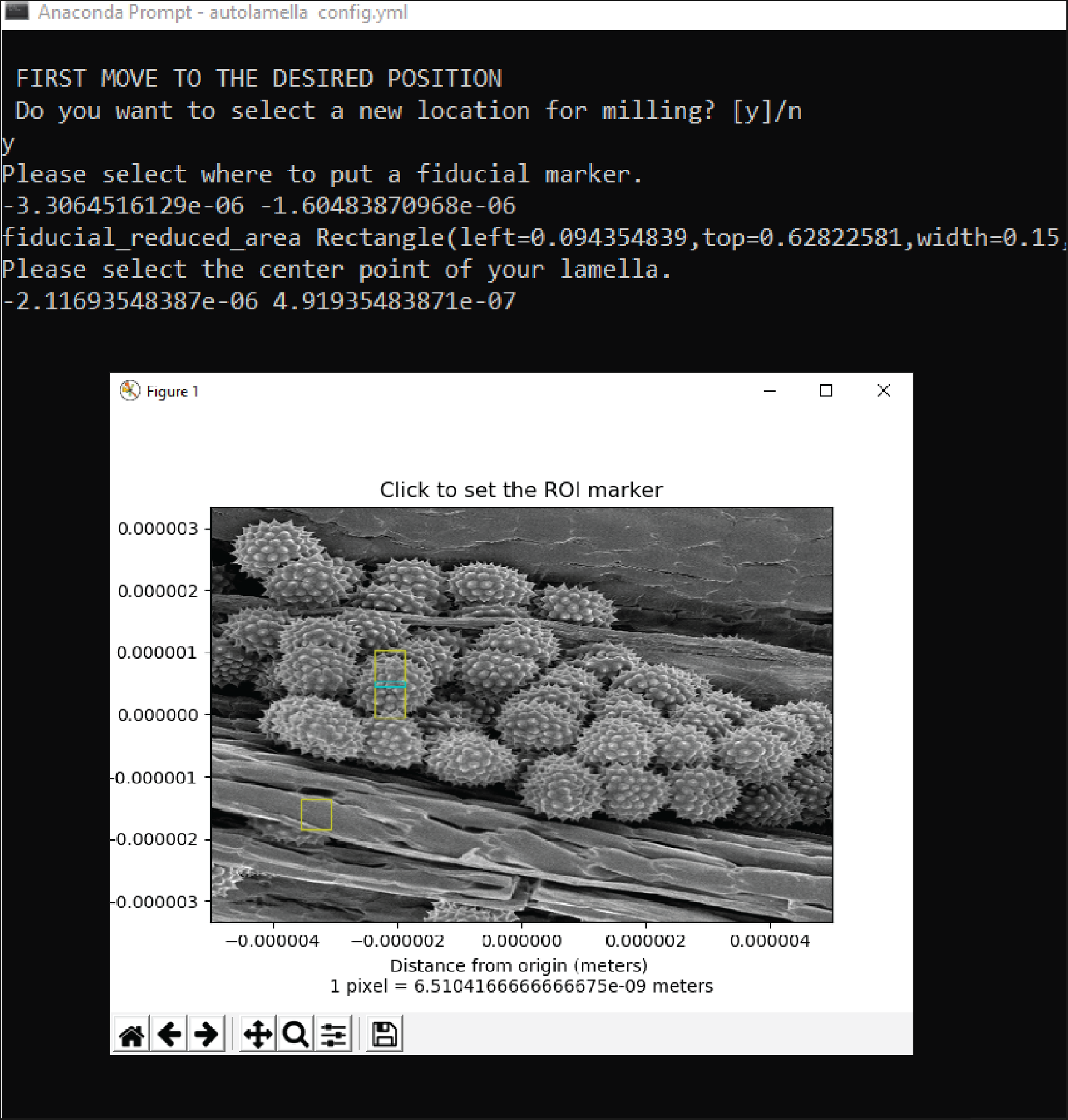
Screenshot of the interactive selection of the lamella position. The command-line prompt asks the user to “Please select the centre point of your lamella” and a new pop up window displays the current ion beam image with the fiducial marker position shown for reference. The user clicks on the image to choose the position for the lamella. The cyan rectangle indicates the position of the final lamella, and the surrounding yellow box indicates the total size of the trenches as defined in the configuration file. The user can click on in multiple locations to optimise the position of the lamella, and the last coordinates chosen before closing the window will be used for milling. Screenshots have been produced using Autoscript simulation mode.

**Supplementary figure 6:**
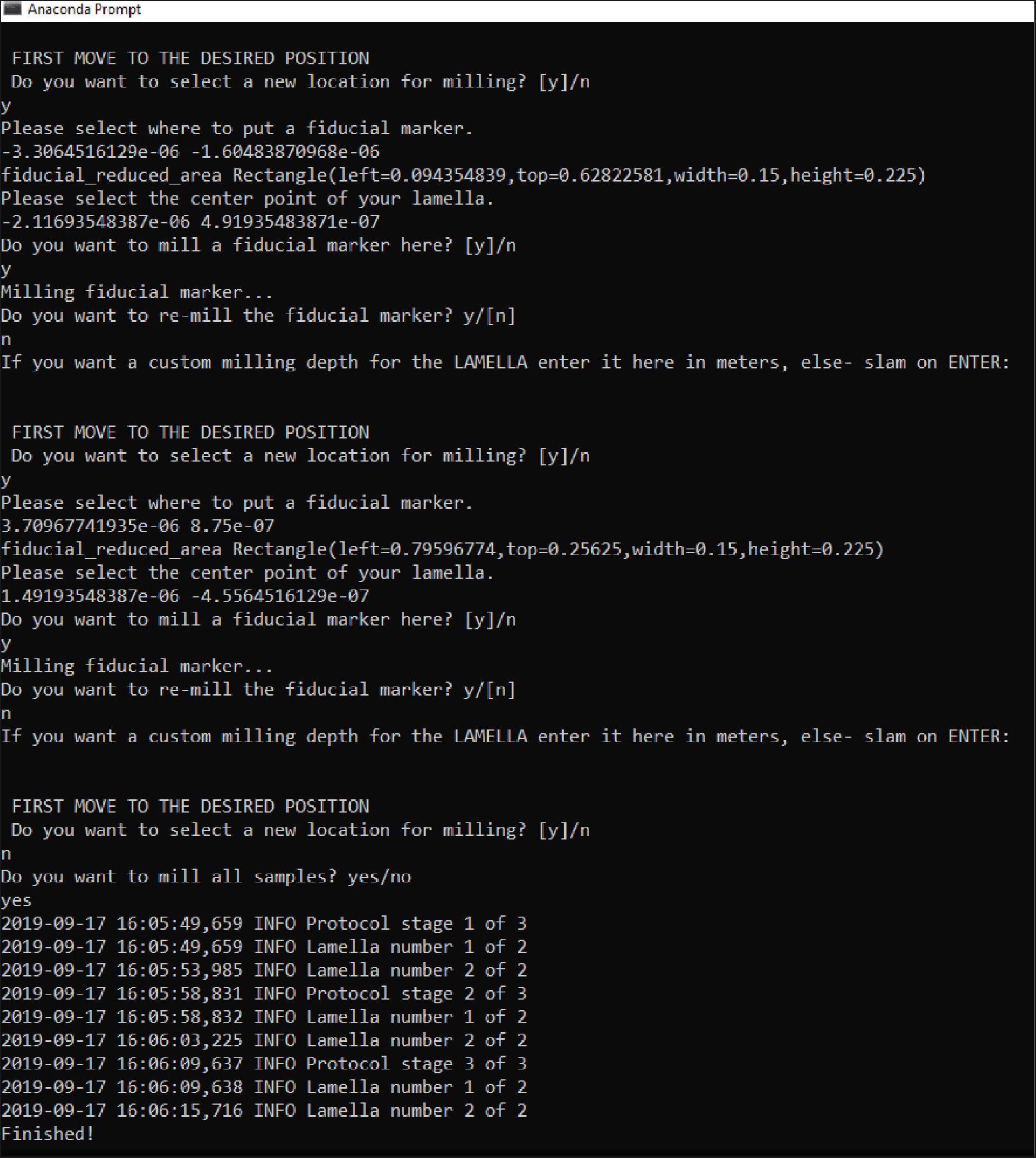
Example output from the command prompt for an automated batch job when preparing two lamellae. After the fiducial mark has been prepared the program images it and provides the option to mill further in case the depth selected does not provide suitable contrast on one particular area. Further the lamella depth can be tuned for each position in case the sample has significant variations. the value should be in meters (e.g. 1e-6 for 1 μm), if the depth specified in the configuration file is acceptable the user should not enter any value. If no more locations are selected (negative answer at the question “Do you want to select a new location for milling?)” the interactive section ends at the question “Do you want to mil all samples?”. This screenshot demonstration has been produced using the Autoscript simulation mode.

**Supplementary video 1: slice by slice video of the tomogram shown in figure 4.**

